# Selective targeting of IL-1RAP-dependent eosinophilic inflammation in allergic fungal airway disease

**DOI:** 10.1101/2025.06.09.658569

**Authors:** Thomas J Williams, James S Griffiths, Luis E Gonzales-Huerta, David Bell, Anna K Reed, Anand Shah, Julian R Naglik, Darius Armstrong-James

## Abstract

It is estimated that in excess of 10 million people around the globe are affected by severe asthma and fungal sensitisation (SAFS) or allergic bronchopulmonary aspergillosis (ABPA), severe asthma endotypes driven by hypersensitivity to environmentally ubiquitous fungal pathogens, primarily *Aspergillus fumigatus*. Here, we sought to define the immunological pathways underlying these allergic fungal airway diseases. To do so, we exploit the chronic exposure repeat challenge model using live *A. fumigatus* conidia to systematically define the key immunological pathways driving airway inflammation in allergic fungal airway disease. In response to daily intranasal challenge, we observed increased absolute numbers of neutrophils and eosinophils in bronchoalveolar lavage fluid (BALF), characteristic of human allergic fungal airway disease, with significant depletion of the alveolar macrophage population. Transcriptomic analysis of BALF cells identified increased expression of IL-1 family cytokines and receptors including IL-1β, IL-1RL1, IL-1R2 and the IL-18 binding protein. Complementary proteomic analysis of BALF revealed increased levels of cell death related proteins calprotectin and IL-1 Receptor Accessory Protein (IL-1RAP). Targeting IL-1RAP, using knockout mice, led to selective reduction in eosinophilia, IL-5 and IL-13 in the airways without impairment of fungal killing. This study identifies a role for IL-1RAP in the generation of eosinophilia independent of neutrophil influx and highlights its potential as a novel immunotherapeutic target for the treatment of allergic fungal airway disease.

**Author Summary:** Allergic bronchopulmonary aspergillosis (ABPA) is a form of lung disease primarily affecting those with asthma or cystic fibrosis induced by the ubiquitous pathogen *Aspergillus fumigatus*. Understanding the immunological mechanisms contributing to this airway disease is important for the development of novel treatments to improve quality of life and preserve lung function. In this study, we show that an *Aspergillus fumigatus* repeat challenge model phenocopies immunological features of allergic fungal airway disease and is hallmarked by increased IL-1 family signalling. We identify the IL-1RAP as a contributor to eosinophilia and the release of Type 2 cytokines during allergic fungal airway disease. This work represents major step in describing the molecular mechanisms of airway eosinophilia in response to fungal exposure and lays the groundwork for further dissection of inflammatory pathways to target with immunotherapeutic approaches.

## Introduction

The opportunistic pathogen *Aspergillus fumigatus (Af)* is found ubiquitously in the air we breathe^1^. Primary clinical presentations of *Aspergillus*-related disease complicating asthma are severe asthma with fungal sensitisation (SAFS) and allergic bronchopulmonary aspergillosis (ABPA)^2^. The combined global burden of SAFS and ABPA is estimated to exceed 10 million people^3,4,5^. Long term ABPA may result in recurrent asthma exacerbations, steroid dependence, the development of bronchiectasis or pulmonary fibrosis, irreversible lung damage and the development of invasive disease^6^.

Commonly, murine models of allergic fungal airway disease are generated through sensitization of the mouse by intraperitoneal injection of *Aspergillus* extract with or without adjuvant^7^. This is followed by intranasal challenge with either *Aspergillus* extract or large doses of live *Af* conidia, however these models do not reflect the continuous human exposure to live conidia^8^. To address this issue models have been developed based on repeat or continuous airway exposure to live conidia^7,9^. These models have the additional benefit that they allow the true impact of immunotherapeutic interventions on both airway inflammation and control of fungal infection to be assessed.

IL-1 family members play an important role in the host response to infectious and immunological challenges. Two of the most well-known members of this family are IL-1α and IL-1β, highly inflammatory cytokines which when excessively released can lead to severe pathological consequences^10^. The cytokine IL-33 is another IL-1 family member that has been implicated in allergic airway inflammation ^11^. Furthermore, fungal pathogens such as *Af* induce diverse programmed cell death responses in several immune cells, leading to the release of biologically active IL-1 family members^12,13^.

The IL-1 receptor family has been implicated in both cell death and allergy due to interactions with IL-1β and IL-33^14^. IL-1 receptor 1 (IL-1R1) forms a ternary complex with IL-1β and IL-1R accessory protein (IL-1RAP, also known as IL-1R3) resulting in NF-κB activation for a pro-inflammatory signal^15,16^. IL-1RAP can also form ternary complexes with other IL-1R family members such as with IL-1 Receptor-like 1 (IL-1RL1, also known as ST2) and IL-33, or IL-1RL2 and IL-36, leading to similar pro-inflammatory signals^17,18^. Given the involvement of these pathways in allergic airway diseases, these observations make it an attractive target for immunotherapy.

Combining transcriptomic and proteomic approaches we identified IL-1 family cytokines and receptors as central mediators of inflammatory signalling during experimental allergic fungal airway disease. Given the development of novel biologics targeting IL-1RAP^19–21^ we therefore investigated the impact of IL-1RAP deletion in this context. This led to selective reduction of eosinophil recruitment to the airway without increased fungal burden. Overall, we highlight a role for IL-1RAP in the eosinophilic response during allergic fungal airway disease suggesting that therapeutics targeting this receptor have potential for treating those with allergic fungal airway disease.

## Methods

### Mouse Ethics and Housing

All mouse experiments were approved by the United Kingdom Home Office and the Imperial College animal ethics committee, performed in accordance with the project licenses PPL PF9534064 and PP6711863. Male C57BL/6J male mice (6-8 weeks old) were ordered from Charles River, UK. B6;129S1-Il1raptm1Roml/J (IL-1RAP^-/-^) were purchased from Jax (RRID:IMSR_JAX:003284) and backcrossed for 10 generations to C57BL/6J. All mice were housed in individually vented cages with free access to autoclaved food and water. Mice were assigned to experimental groups at random and, where possible, mixed among cages.

### Aspergillus culture

*Aspergillus fumigatus* strain CEA10 (FGSC A1163) was obtained from the Fungal Genetics Stock Centre, an eGFP-expressing strain (ATCC46645-eGFP) was kindly gifted from Frank Ebel. All strains were cultured on Sabouraud dextrose agar (Oxoid). Conidia were harvested in 0.1% Tween/H2O and filtered through MIRACLOTH (Calbiochem, UK).

### Repeat Challenge Aspergillus Model

Using the approach employed by Porter et al.^9^, mice were intranasally dosed with 2×10^5^ resting conidia in 50μl of PBS while under isoflurane anaesthesia. Doses were repeated for 3, 7 and 14 days with mice being culled either 3 or 24 hours after the final dose by intra-peritoneal overdose of pentobarbital.

### Histology

Lungs were inflated, excised and fixed in 10% neutral buffered formalin overnight and moved into 70% ethanol. Lungs were then wax embedded, sectioned to 4μm sections and H&E or PAS stained. %Inflammation and %Mucus were determined by threshold image analysis, using Image J software.

### Bronchoalveolar Lavage

Bronchoalveolar lavage (BAL) was performed by tracheal cut-down and intubation with a self-designed catheter. Lungs were lavaged by 3 separate instillations of 1ml of PBS/2mM EDTA. The supernatant fraction of the first instillation was used for LDH, Bradford, ELISA and Proteomics. The cell fractions of all 3 instillations were combined and used for flow cytometry. For CFUs all instillations were combined.

### Colony Forming Unit Assay

Serial dilution of BAL fluid was performed and plated onto Sabouraud dextrose agar plates in triplicate. Plates were incubated for 18-24 hrs at 37°C and CFUs were manually counted.

### Flow Cytometry

Cell pellets were stained according to the antibody details in Table 2.2 below, fixed in 2% formaldehyde (28908, ThermoFisher Scientific) and resuspended in FACS buffer (PBS, 1% FBS, 3mM EDTA). 123count eBeads counting beads (01-1234-42, ThermoFisher Scientific) were used for cell enumeration. For determination of GFP+ *Af* cells, CEA10 infected mice were used as threshold. Flow Cytometry was performed on an LSR Fortessa III flow cytometer (BD Biosciences). Data were analysed using FlowJo (Treestar).

| Target Species | Target Protein | Fluorophore | Clone | Company | Cat. Number |
| --- | --- | --- | --- | --- | --- |
| Mouse | Live/Dead | Blue | N/A | Thermo Fisher Scientific | L34959 |
|  | CD19 | BV785 | 6D5 | Biolegend | 115543 |
|  | CD45.2 | BUV737 | 104 | BD | 564880 |
|  | CD3 | BUV395 | 17A2 | BD | 740268 |
|  | Ly6C | PE Cy7 | HK1.4 | eBioscience | 25-5932-82 |
|  | MHC-II-I/A I/E | APC Cy7 | M5/114.15.2 | Biolegend | 107628 |
|  | Ly6G | AF700 | 1A8 | Biolegend | 127622 |
|  | CD11b | BV605 | M1/70 | Biolegend | 101237 |
|  | F4/80 | APC | BM8 | Invitrogen | 17-4801-82 |
|  | CD11c | PE-Cy7 | HL3 | BD | 558079 |
|  | CD11c | FITC | HL3 | BD | 557400 |
|  | Siglec F | PE | E50-2440 | BD | 562068 |

### Bradford Assay

Basic protein quantification of BAL supernatant was carried out using Pierce Coomassie Plus (Bradford) Assay Reagent (23238, ThermoFisher Scientific), concentrations were calculated based on a BSA standard curve.

### Transcriptomics

Bulk RNA was isolated from total bronchoalveolar lavage cells using QIAshredder (79654, Qiagen) and RNeasy Mini Kits (74104, Qiagen) according to the manufacturer’s instructions. RNA quantity and purity were assessed by Nanodrop. Sample integrity was checked using a BioAnlayzer (Agilent) Library preparation was carried out by Novogene (Novogene (UK) Ltd). Messenger RNA was purified from total RNA using poly-T oligo-attached magnetic beads. After fragmentation, the first strand cDNA was synthesised using random hexamer primers, followed by the second strand cDNA synthesis using dTTP. The library was checked with Qubit and real-time PCR for quantification and Bioanlayzer for size distribution detection. Quantified libraries were pooled and sequenced on Illumina platforms.

Raw data (raw reads) of fastq format were firstly processed through fastp software. Clean data (clean reads) was obtained by removing reads containing adapter, reads containing ploy-N and low quality reads from raw data. Q20, Q30 and GC content of the clean data was calculated and all downstream analyses were based on the clean data with high quality.

Reads were mapped to the murine genome (GRCm39/mm39, GCF_000001635.27). An index of the reference genome was built using Hisat2 v2.05 and paired-end clean reads were aligned to the reference genome using Hisat v2.05. The mapped reads of each sample were assembled by StringTIe (v1.3.3b) in a reference-based approach.

Analysis of RNAseq libraries was carried out using the Qlucore Multiomics Explorer 3.9 (Qlucore, Sweden). This included determination of differentially expressed genes, volcano plot and heatmap generation. Genes with an adjusted P-value <=0.05 and fold change >2 were assigned as differentially expressed. Gene Ontology (GO) enrichment analysis – GO enrichment analysis was implemented by the clusterProfiler R package. GO terms with a corrected P-value <=0.05 were considered significantly enriched by differentially expressed genes. The RNA-seq data generated during this study are available at NCBI Sequence Read Archive: PRJNA1250534.

### Proteomics

BAL supernatant was taken and spun down at 16000 g for 1 min in a 0.22 μm filter containing centrifuge tube. Undesirable proteins were removed from the samples using a Multiple Affinity Removal Spin Cartridge Mouse 3 (5188-5289, Agilent) according to the manufacturer’s instruction. Samples then underwent acetone precipitation, by addition of ice-cold acetone at a 1:5 ratio, at –80C for 1 hour. Precipitate pellets were dried, resuspended in a buffer containing 10mM TCEP, 40Mm Chloroacetamide, 1%SDC, 100mM tris-HCL P.H 8.8 and placed on a heat block at 95C for 10 mins. 0.5 μg Trypsin/LysC was added to the samples and left to digest overnight at room temperature, acetonitrile and TFA were added for final concentrations of 6% and 0.6% respectively. Samples were stored at −80 °C until analysis.

All samples were injected and separated on a 1290 liquid chromatography system (Agilent) operating in normal flow mode at 200 μl/minute. 10-20 μl of the sample was separated on a AdvanceBio Peptide Plus column (2.7 μm particle size, 2.1 i.d. x 50mm (Agilent)). A 19.5-minute method with the following gradient was used: 97% Buffer A (0.1% (v/v) formic acid in water), 3% Buffer B (0.1% (v/v) formic acid in acetonitrile). Buffer B was increased to 40% over 12.5 minutes, followed by an increase to 100% buffer B in 2 minutes, where it was held for 2 minutes. Buffer B was then ramped back down to 3% in 1 minute and equilibrated for 2 minutes prior to the next injection. The 1290 LC system was coupled to a 6550 iFunnel Q-ToF Mass Spectrometer (Agilent) equipped with an iFunnel electrospray source and running MassHunter Data Acquisition software.

For relative peptide abundance, data was acquired in MS only mode over the m/z range 300-1700 at the rate of 1 spectrum per second. Peptides were identified using an auto-MS/MS method selecting up to 20 precursors per cycle, with a threshold of 5000 counts and a charge state of 2 or greater. Profinder B10 (Agilent) was used for peak list generation. The resulting MS only files were imported into Mass Profiler Professional (Agilent) for relative peptide abundance comparison. Samples were grouped according to type and features (as defined by accurate mass and retention time) were clustered for further analysis. Results were filtered such that a feature was present in all samples within both groups. Log/log of the intensity of features in each sample group was plotted and a paired t-test (p <0.05) was carried out to identify features that had a fold change of 2 or higher.

LC/MS/MS data was extracted into peak lists using SpectrumMill (Agilent). Database searching against SwissProt was achieved using the following search parameters; enzyme specificity: trypsin, charge states: 2 or greater, fixed modification: Cysteine carbaminoacetylation, and variable modifications: Methionine oxidised. A score of 11 was set for acceptance of peptide assignments and 19 for protein identifications as well as %SPI (the percentage of the extracted spectrum that is explained by the database search result) (>60%).

Analysis of proteomic results was carried out using Qlucore Multiomics Explorer 3.9 (Qlucore, Sweden). This included determination of significantly different proteins, volcano plot and heatmap generation. Proteins with an adjusted P-value <=0.05 were assigned as significant.

Functional protein association networks creation for the identified significant proteins was created using the STRING database and online platform (string-db.org). Clusters were created using MCL clustering (Inflation score >4).

### Lactate Dehydrogenase (LDH) Assay

LDH levels in BAL supernatants were measured using the CytoTox 96® Non-Radioactive Cytotoxicity Assay kit (Promega) according to the manufacturer’s instructions.

### ELISA

ELISAs were performed according to the manufacturer’s instruction using the murine TNF-a, CXCL1, IL-1β, IL-33, IL-5 and IL-13 DuoSet kits from R&D systems.

### Statistics

All data are expressed as mean ± SEM unless otherwise stated. To assess statistical significance, one-way ANOVA (≥3 groups), two-way ANOVA and Students t-test/Mann-Whitney test (2 groups) were carried out. For all figures, p values are represented as followed: *P<0.05, **P<0.01, ***P<0.001, ****P<0.0001. Statistical analyses were carried out using GraphPad Prism (La Jolla, CA, USA) or Qlucore Multiomics Explorer 3.9 (Qlucore, Sweden).

## Results

### Chronic exposure to Af leads to increased inflammation, mucus production and fungal persistence in the lung

To elucidate the mechanisms of *A. fumigatus* eosinophilia in lung disease we first sought to establish repeat challenge models of allergic fungal airway disease for immune cell, transcriptomic and proteomic investigations. Using the approach employed by Porter *et al.*^9^, C57BL/6J were dosed intranasally for 14 days with 2×10^5^ resting *Af* conidia (Fig 1A). Mice were monitored for mortality and assessed for any signs of morbidity including weight loss, lethargy, and hunching. After 14 days of dosing the mortality rate was zero (Fig 1B), and the *Aspergillus* exposed mice did not lose significant amounts of weight (Fig 1C). H&E staining demonstrated cellular infiltrates and inflammation of the airways of *Af* exposed mice (Fig 1D), which was significantly increased on thresholding quantification (Fig 1E). *Af* infected mice also exhibited mucus production that was not present in the PBS controls (Fig 1F and G). On histological staining there was no clear evidence of *Af* conidia or hyphae present in the lungs, however CFU counts on bronchoalveolar lavage fluid (BALF) revealed low levels of *Af* in the airways (Fig 1H), suggesting that whilst the majority of conidia are killed there is a low-level persistence of fungus in the airway.

**Figure 1:**
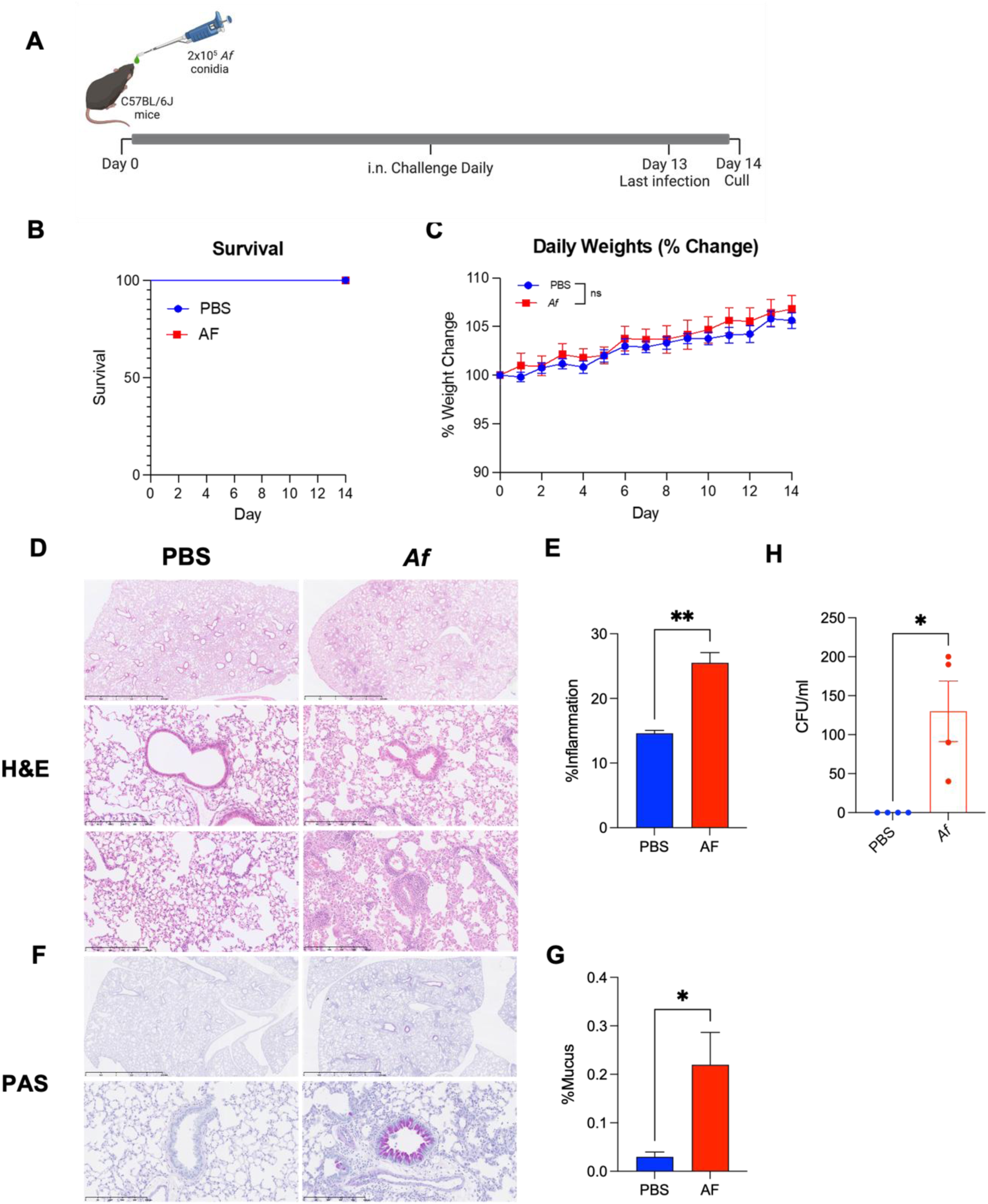
Increased inflammation, mucus production and fungal persistence in the lung of mice with allergic fungal airway disease. **A)** Mice were dosed daily with 2×10^5^ live *A. fumigatus* conidia or PBS for 14 days and culled 24 hours after the final dose. Mice were monitored for **B)** survival and **C)** weight change (n=9). **D-G)** Representative lung sections stained for **D)** H&E and **F)** PAS. **E)** %Inflammation and **G)** % Mucus was determined by threshold image analysis. (N=3) **H)** Bronchoalveolar lavage fluid was collected and CFUs were counted (N=4). Data represents mean ± SEM from at least two independent experiments. C) Two-way ANOVA, E, G, H) Students T-test, *P<0.05, **p<0.01. CFU – Colony forming unit, H&E – Hematoxylin & eosin, PAS – Periodic acid Schiff.

### Chronic exposure to Af leads to eosinophilia and neutrophilia in the airways

We next sought to establish which cellular populations were present in the airways and how they changed over time in allergic fungal airway disease. Mice were culled 24 hours after having 3, 7 or 14 intranasal doses of *Af* and the cells in the BAL fluid were analysed by flow cytometry (Fig 2A, Fig S1). After 7 days of exposure, eosinophils represented ∼40% of airway immune cells, increasing to ∼75% after 14 days (Fig 2B, Fig S2A). Significant influx of neutrophils occurred after 3 doses of *Af* (Fig 2C, Fig S2B). After 14 days of exposure, a significant reduction in the number of alveolar macrophages was observed (Fig 2D, Fig S2C). A significant infiltration of monocytes is seen after 3 days of exposure, increasing by day 7 (Fig 2E, Fig S2D). We observed significant increases in the number of T cells after 7 days of exposure which was increased again by day 14 (Fig 2F, Fig S2E), while B cells were significantly increased after 14 days (Fig 2G, Fig S2F).

**Figure 2:**
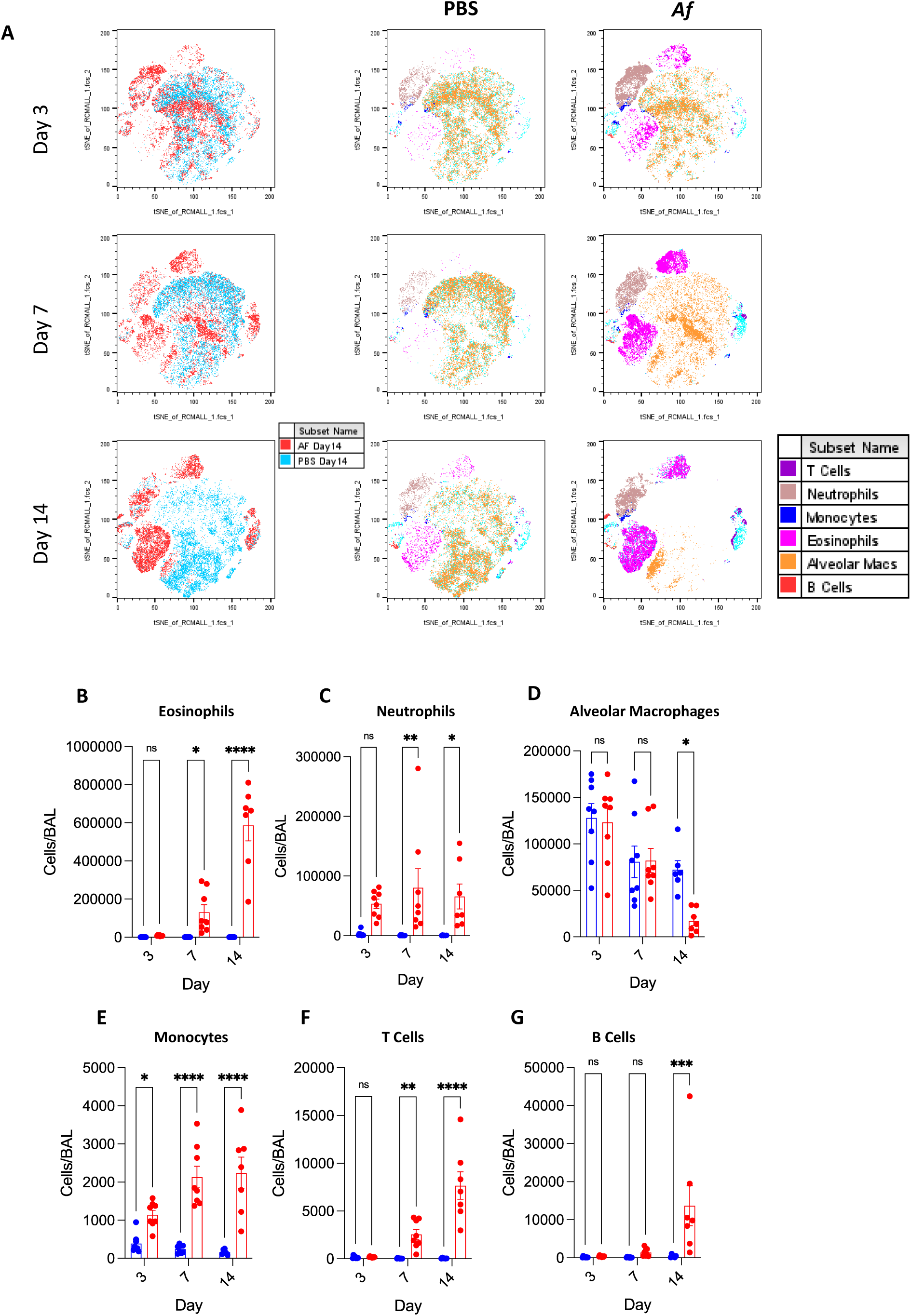
The cellular compartment of the airways changes significantly over the course of chronic exposure to *A. fumigatus.* Mice were dosed daily with 2×10^5^ live *A. fumigatus* conidia or PBS for up to 14 days and culled 24 hours after the final dose Flow cytometry was carried out to determine the immune cell populations in the airways. **A)** tsne plots representing the airway immune cell composition at 3, 7 and 14 days of *A. fumigatus* exposure. Enumeration of immune cells at 3, 7 and 14 days of *A. fumigatus* exposure for **B)** Eosinophils, **C)** Neutrophils, **D)** Alveolar Macrophages, **E)** Monocytes, **F)** T Cells and **G)** B Cells (N=4-8). Data represents mean ± SEM from at least two independent experiments. B-G) One-way ANOVA, *P<0.05, **P<0.01, ***P<0.001, ****P<0.0001.

Approximately 1% of alveolar macrophages and 1.5% of neutrophils were in direct contact with *Af* conidia, highlighting their role as first lines of defence and primary phagocytes. We also observed a very small population, 0.05%, of eosinophils interacting directly with the conidia (Fig S2G and H).

### Chronic exposure to Af leads to cell death and increased levels IL-1 family cytokines and receptors

To systematically characterise the immune response during repeated *Aspergillus* exposure, we next carried out bulk transcriptomic analysis of the BAL cells. We identified 443 differentially expressed genes, of which 256 were upregulated and 187 were downregulated in mice exposed to *Af* for 14 days compared to PBS controls (Fig 3A and B, Table S1 and S2). Gene Ontology (GO) enrichment analysis showed upregulation of pathways expected during the generation of allergy including granulocyte migration, T cell activation and differentiation, and myeloid cell proliferation (Fig 3C, Table S3), while downregulated pathways included autophagic pathways and nitric oxide biosynthesis/metabolism (Fig 3D, Table S3). Investigating the differentially expressed genes at an individual level revealed several upregulated genes encoding for interleukins and interleukin receptors. These included the Type 2 cytokines IL-4 and IL-13, as well as receptors including IL-2 receptor beta unit (IL-2rb), IL-21 receptor (IL21r), IL-27 receptor alpha subunit (IL-27ra) and IL-20 receptor beta unit (IL-21rb). Genes encoding the IL-1 family signalling members IL-1 receptor-like 1 (IL-1rl1/ST2), IL-1 receptor 2 (IL-1r2), IL-1β and IL-18 binding protein (IL-18bp) were also significantly upregulated during repeat fungal challenge (Fig 3E).

**Figure 3:**
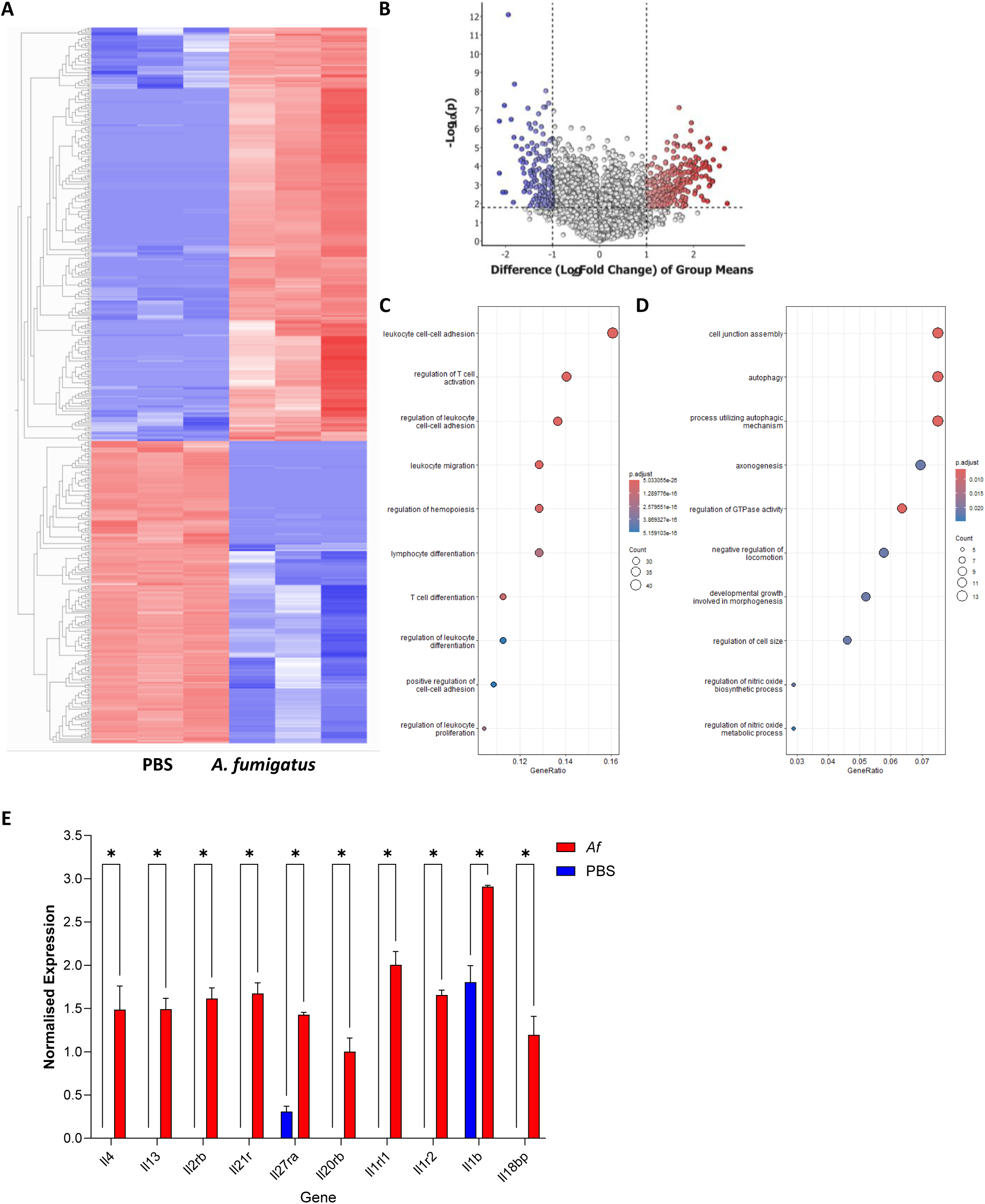
Increased damage markers and inflammatory cytokines in the airways of mice with allergic fungal airway disease. Mice were dosed daily with 2×10^5^ live *A. fumigatus* conidia or PBS for 14 days and culled 24 hours after the final dose. Bulk RNAseq was carried out on bronchoalveolar lavage cells. **A)** Heatmap and **B)** showing all differentially expressed genes (determined as padj>0.05 and fold-change>2) compared to PBS controls. curated list of differentially expressed genes of interest. Dot plot of GO pathway analysis representing the top 10 enriched pathways for **C)** upregulated and **D)** downregulated differentially expressed genes in allergic fungal airway disease **E)** Bar graph showing normalized expression of upregulated interleukins and interleukin receptors in allergic fungal airway disease. E) Students T-test, *P<0.05

Having observed increased expression of IL-1 family cytokines and receptors we further characterised the BALF to investigate the amount of inflammatory cell death occurring in the airway. As a proxy of inflammatory cell death, we measured the levels of lactate dehydrogenase (LDH) and free protein in the airways. At 14 days there was a significant increase in the levels of both LDH and free protein in the exposed mice (Fig 4A and B). To complement our transcriptomic results, we undertook proteomic analysis of the BALF identifying 164 proteins. Of these 164 proteins, 42 were differentially expressed, 38 increased and 4 decreased, in *Af treated* mice compared to PBS controls (Fig 4C and D) (Table S4). These proteins included the classic DAMP calprotectin (S100A8/S100A9) and the IL-1 family signalling co-receptor IL-1 receptor accessory protein (IL-1RAP) as well as complement proteins, chitinases and pulmonary surfactants. Having further evidence of IL-1 family signalling by the identification of IL-1RAP, we examined the levels of the IL-1 family cytokines that signal through IL-1RAP, IL-1β and IL-33. We observed significant increases in the levels of both IL-1β and IL-33 in the BALF of mice exposed to *Af* for 14 days (Fig 4E-G). This, taken together with the increased levels of LDH and upregulation of IL-1 family receptors, suggests that inflammatory cell death is occurring in the airways of mice with allergic fungal airway disease followed by the release and subsequent signalling of IL-1 cytokines through receptor complexes containing IL-1RAP.

**Figure 4:**
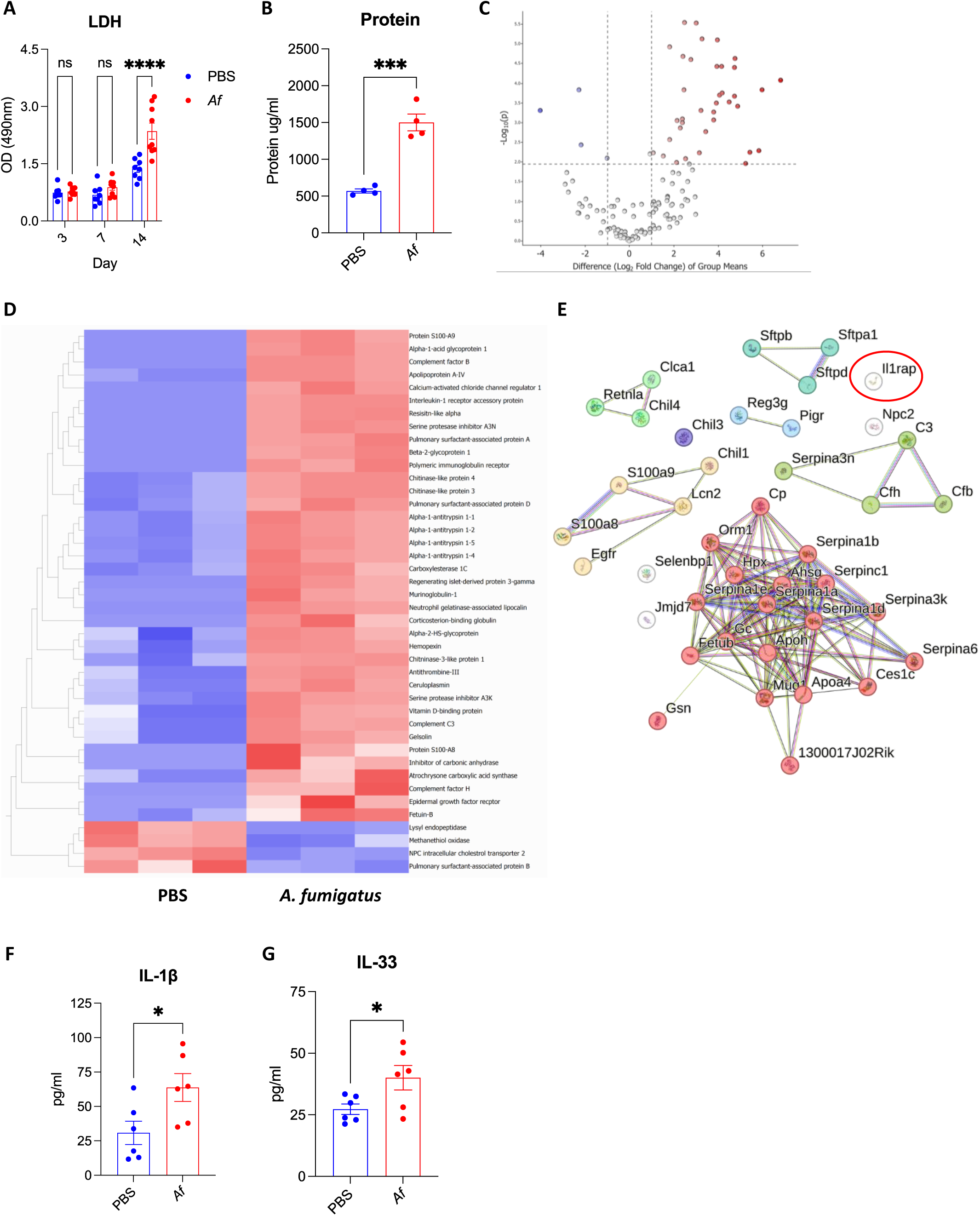
Proteomic analysis reveals increased IL-1RAP, IL-1β and IL-33 in allergic fungal airway disease. Mice were dosed daily with 2×10^5^ live *A. fumigatus* conidia or PBS for 14 days and culled 24 hours after the final dose. Bronchoalveolar lavage fluid was assessed for **A)** LDH (N=8-9) and **B)** Total protein (N=4). **C and D)** LC-MS/MS analysis was carried out on bronchoalveolar lavage fluid to identify differentially expressed proteins (padj>0.05, fold change>2) (N=3) **C)** volcano plot of all identified proteins, **D)** Heat-map of differentially expressed proteins. E) Functional protein network association created using STRING software for all 42 differentially expressed proteins, clustered using MCL clustering (>4). Bronchoalveolar lavage fluid was assessed by ELISA for **F)** IL-1β and **G)** IL-33 (N=6). Data represents mean ± SEM from at least two independent experiments. A) One-way ANOVA, B, F, G) Students T-test, *P<0.05, ***P<0.001, ****P<0.0001.

IL-1RAP is a key co-receptor in the IL-1 family signalling pathway, acting as a co-receptor for IL-1r1 which binds IL-1α and IL-1β, for IL-1rl1 (ST2) which binds IL-33 and for IL-1rl2 (IL-36r) which binds IL-36, making it essential for effective IL-1 family signalling. Having identified increased levels of IL-1β and IL-33, upregulation of IL-1rL1 and increased levels of IL-1RAP in the airways, we next sought to investigate the role that IL-1RAP plays in the generation of allergic fungal airway disease using IL-1RAP knockout mice (IL-1RAP^-/-^). No mortality or significant weight loss were observed in the IL-1RAP^-/-^ mice compared to wildtype (WT) controls during allergic fungal airway disease (Fig 5A). We observed a significant reduction in the number of eosinophils in the airways of IL-1RAP^-/-^ mice compared to WT controls (Fig 5B). However, no significant difference was observed in the number of alveolar macrophages or neutrophils (Fig 5C and D). Additionally, we found no significant difference in fungal control between IL-1RAP^-/-^ mice and WT controls (Fig 5E). Finally, having observed reduction in the eosinophil numbers in the airway, we measured levels of IL-33, IL-5 and IL-13, cytokines typically associated with the allergic response and eosinophil chemotaxis. We did not observe any difference in airway levels of IL-33 (Fig 5F), however there were significant reductions in the amounts of both IL-5 and IL-13 (Fig 5G and H). This suggests that deficiency in IL-1RAP results in reduced eosinophilia due to reduced levels of the downstream eosinophil chemoattractants IL-5 and IL-13, likely as a result of the inability to signal through the IL-33/IL-1RL1/IL-1RAP signalling axis.

**Figure 5:**
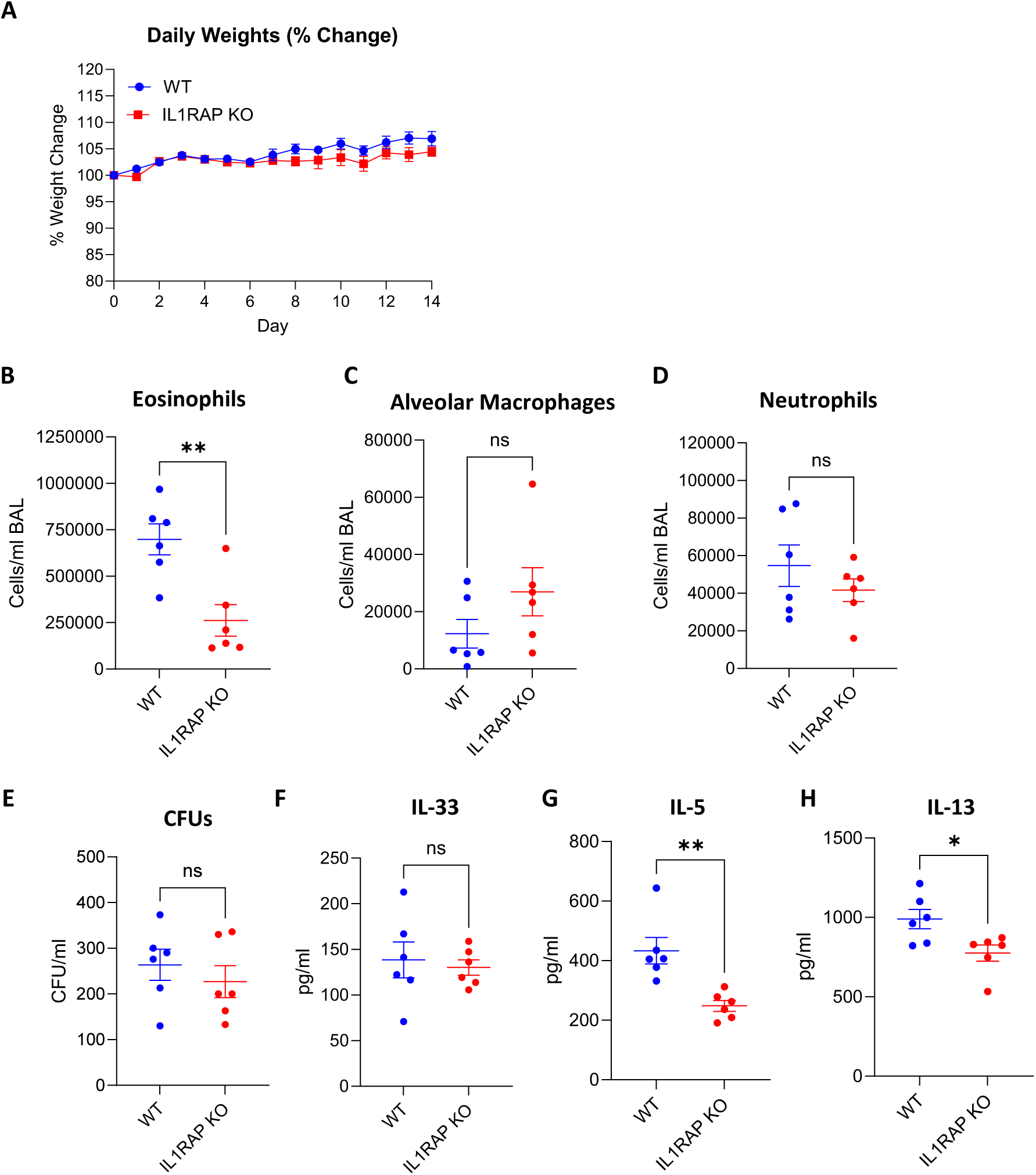
IL-1RAP deficiency reduces eosinophilia during chronic exposure to *A. fumigatus* without loss of fungal control. IL1RAP^-/-^ and WT mice were dosed daily with or without 2×10^5^ live *A. fumigatus* conidia for 14 days and culled 24 hours after the final dose. Mice were monitored for **A)** weight change. **B-H)** Bronchoalveolar lavage fluid (BALF) was collected and assessed for **B)** eosinophil, **C)** alveolar macrophage and **D)** neutrophil counts, **E)** CFUs and levels of **F)** IL-33, **G)** IL-5 and **H)** IL-13 (n=6). Data represents mean ± SEM from at least two independent experiments. A) Two-way ANOVA B-H) Students T-test, *P<0.05, **P<0.01.

## Discussion

ABPA is characterised by recurrent inflammatory exacerbations leading to lung damage and eventual decline. Here, we present a systematic characterisation of the inflammatory response in the murine model of allergic fungal airway disease through a multi-omic approach. Using knockout animals, we investigated the role that IL-1RAP plays in the generation of allergic fungal airway disease.

Allergic fungal airway disease is typically characterised by airway eosinophilia, increased IgE levels, increased T cell responses and inflammation of the airways^22^. We observe that the repeat challenge model of allergic fungal airway disease phenocopies immunological features of human allergic fungal airway disease such as sustained eosinophil and neutrophil influx, alveolar macrophage loss. These observations were associated with increased levels of cell death and damage markers. Although the source of these markers has not yet been characterised, they are likely a combination of damaged epithelia, dying alveolar macrophages and extracellular trap releasing granulocytes.

*Af* is known to cause significant damage to the epithelium during infection, resulting in the release of inflammatory mediators including IL-1α, IL-33, HMGB1, and TSLP^23–28^. In this model, repeat dosing of *Af* conidia represents a constant assault on the epithelium likely leading to significant epithelial damage. The loss of alveolar macrophages together with increased LDH and IL-1β levels is also indicative of cell death. While *Af* can lyse cells, through the process of germination^29^, the lack of any evidence of fungal hyphae by histology suggests that fungal spores are being controlled. Therefore, any *Af*-induced death is likely to be a result of programmed cell death pathways, for which *Af* can induce a broad range of types^12,30,31^. Extracellular trap release from neutrophils and eosinophils may also contribute the inflammatory profile seen in this model. Neutrophil extracellular traps (NETs) are known to be released in response to *Af* while eosinophil extracellular traps (EETs) have been shown to be present in the sputum of those with ABPA^32–35^. While these ETs restrict hyphal growth, they do not contribute to killing of the fungus and their unrestricted release is considered pathogenic^36–38^. Transcriptomics of airway cells corroborated with plate-based assays identified increased expression and levels of cell death and damage markers including LDH, IL-1α, IL-1β, and IL-33, as well as upregulation of IL-1 family receptor IL-1RL1, in the airways of mice with allergic fungal airway disease.

Proteomic analysis of BAL fluid showed significantly increased levels of chitinase-3 like protein 1 (CHI3L1) and acidic mammalian chitinase (AMCase). CHI3L1 is a known contributor to allergy by contributing to M2 macrophage polarisation and Th2 induction in both skin and food allergies, with polymorphisms in CHI3L1 associated with the development of bronchial asthma and airway remodelling^39–41^. AMCase has demonstrated antifungal properties, targeting of chitin in the fungal cell wall^42,43^. Studies have shown that chitin induces the recruitment of IL-4 expressing immune cells to the lung facilitating allergic inflammation, this was abrogated with exogenous AMCase treatment or overexpression of AMCase in a murine model^44,45^. AMCase has also been implicated in the pathology of allergy^43,46^, it was shown that neutralisation of AMCase ameliorated Th2 dependent inflammation and airway hyperreactivity by inhibiting the IL-13 response^47^.

Moreover, the proteomic analysis of the BAL from mice with allergic fungal airway disease also supports ongoing cell death in the airway with the detection of S100A8 and S100A9 which form the heterodimer calprotectin, a well-established DAMP related to inflammatory cell death and a common marker of tissue inflammation^48–51^. Lastly and of interest, along with increases in the IL-1 family members IL-33 and IL-1RL1 receptor, IL-1RAP was found to be significantly increased in the airways of mice with allergic fungal airway disease. IL-1RAP is an essential co-receptor for IL-1 family signalling, in this context primarily for IL-1β and IL-33 signalling and therefore represents an attractive target for immunotherapeutic intervention. Of note, increased expression of IL-1RAP and IL-33 activation have previously been associated with severe neutrophilic asthma^52^.

Similar to previous studies investigating the role of IL-33 signalling in allergy, in which IL-1RL1 (ST2) knockout mice showed reduced eosinophilia during allergen challenge^53^ IL-1RAP^-/-^ mice with allergic fungal airway disease demonstrated selective reduction of eosinophilia and reduced production of IL-5 and IL-13. The mechanistic basis for these observations is further delineated by the observation that IL-33 challenged IL-1RAP^-/-^ mice also showed reduced eosinophilia and IL-5^54^. The primary producers of these cytokines during allergy are Th2, ILC2 and mast cells signalling through the IL-1RL1/IL-33 axis^55–57^.

Taken together, these observations suggest that IL-1RAP is required for IL-33/IL-1RL1-dependent signalling, IL-5 and IL-13 production and subsequently eosinophilia in allergic fungal airway disease. Monoclonal antibodies targeting IL-1RAP as a pan-IL1 family blockade therapy have been shown to reduce inflammation in murine models of sterile inflammation and are currently undergoing clinical trials^19–21^. Our observations identify IL-1RAP as a novel immunotherapeutically-tractable inflammatory mediator that has potential for selective targeting of eosinophilic inflammation in ABPA.

## Supplementary Material

**Figure S1:**
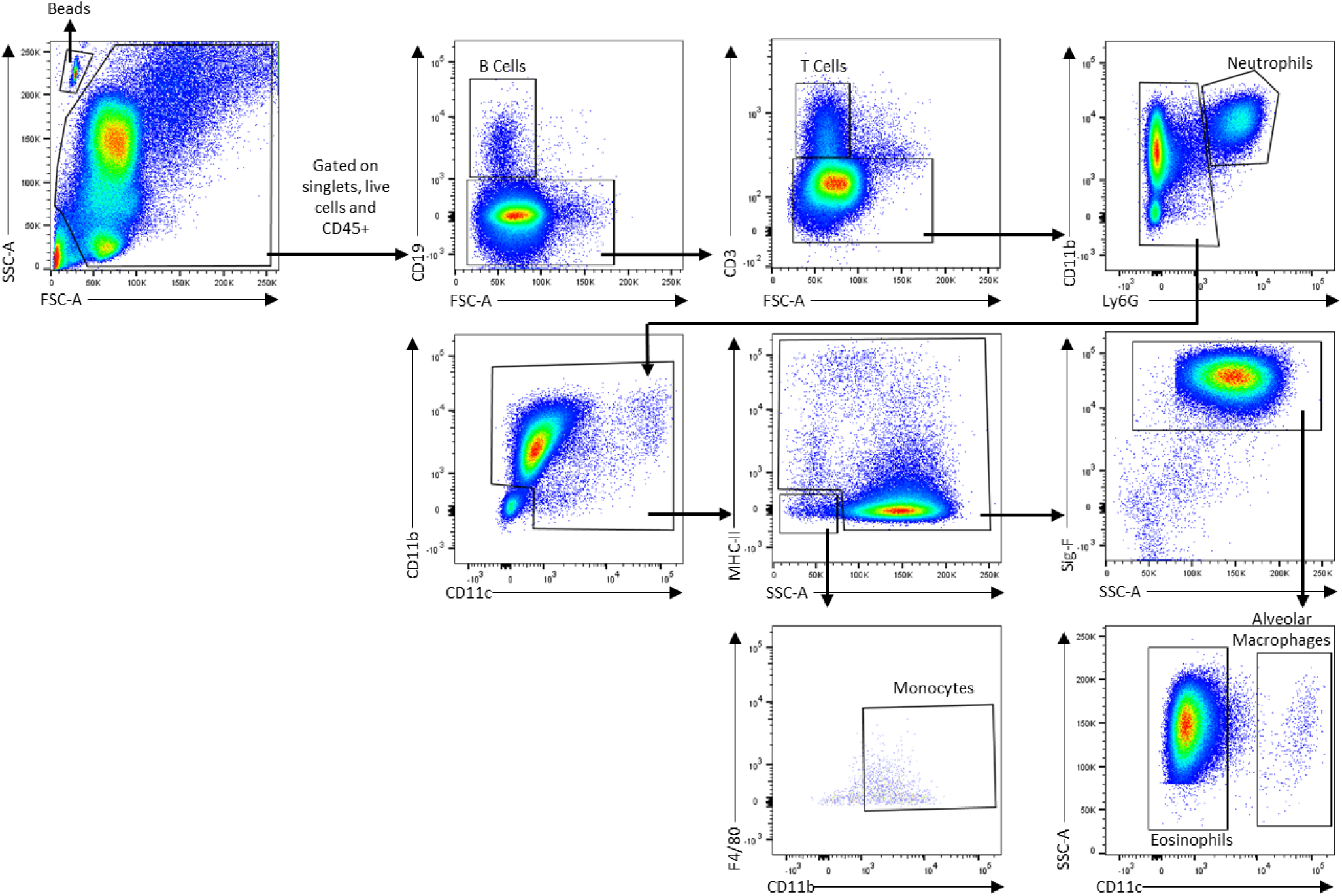
Representative Gating Strategy for immune profiling of murine BAL. Cells were first gated on forward and sidescatter followed by the exclusion of doublets. Dead cells were removed from analysis according to a LIVE/Dead stain and immune cells were identified as CD45+, cell types were subsequently determined through positive and negative getting of cell specific markers. Cell populations were enumerated using flow cytometry counting beads, identified by forward and side scattered, followed by autofluorescence.

**Figure S2:**
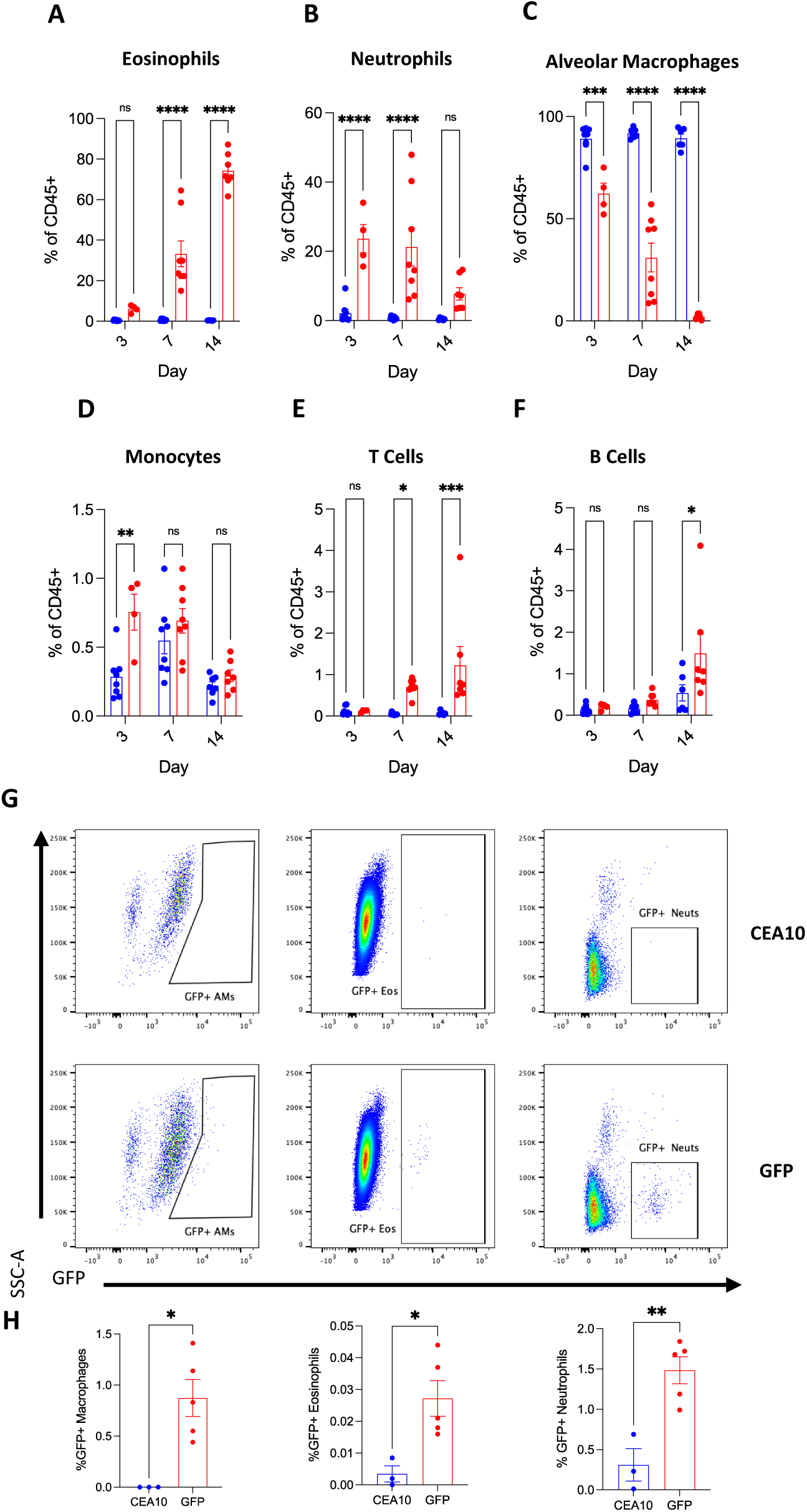
The cellular compartment of the airways changes significantly over the course of chronic exposure to *A. fumigatus.* Mice are dosed daily with 2×10^5^ live *A. fumigatus* conidia or PBS for up to 14 days and culled 24 hours after the final dose Flow cytometry was carried out to determine the immune cell populations in the airways. Proportion of immune cells at 3, 7 and 14 days of *A. fumigatus* exposure for A) Eosinophils, **B)** Neutrophils, **C)** Alveolar Macrophages, **D)** Monocytes, **E)** T Cells and **F)** B Cells (N=4-8). Mice were dosed daily with 2×10^5^ GFP or CEA10 *A. fumigatus* conidia and culled 3 hours after the final dose. Flow cytometry was carried out to assess the fate of GFP expressing *A. fumigatus*. **G)** Representative plots and **H)** Percentage of GFP+ cells in each population for GFP+ Alveolar macrophages, Eosinophils and Neutrophils (N=3-5). Data represents mean ± SEM from at least two independent experiments. A-F) One-way ANOVA, H) Students T-test, *P<0.05, **P<0.01, ***P<0.001, ****P<0.0001.

**Table S1:**
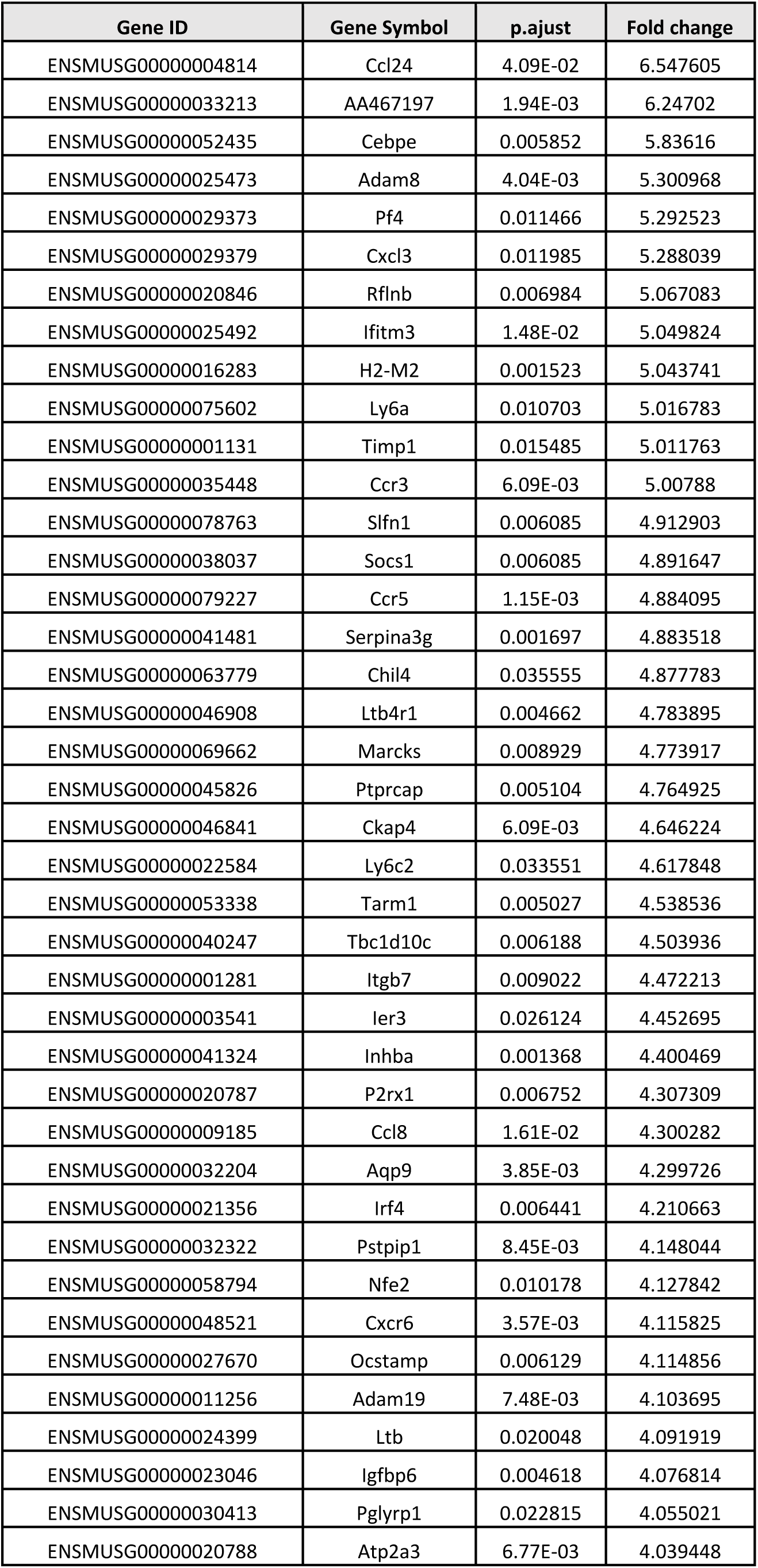
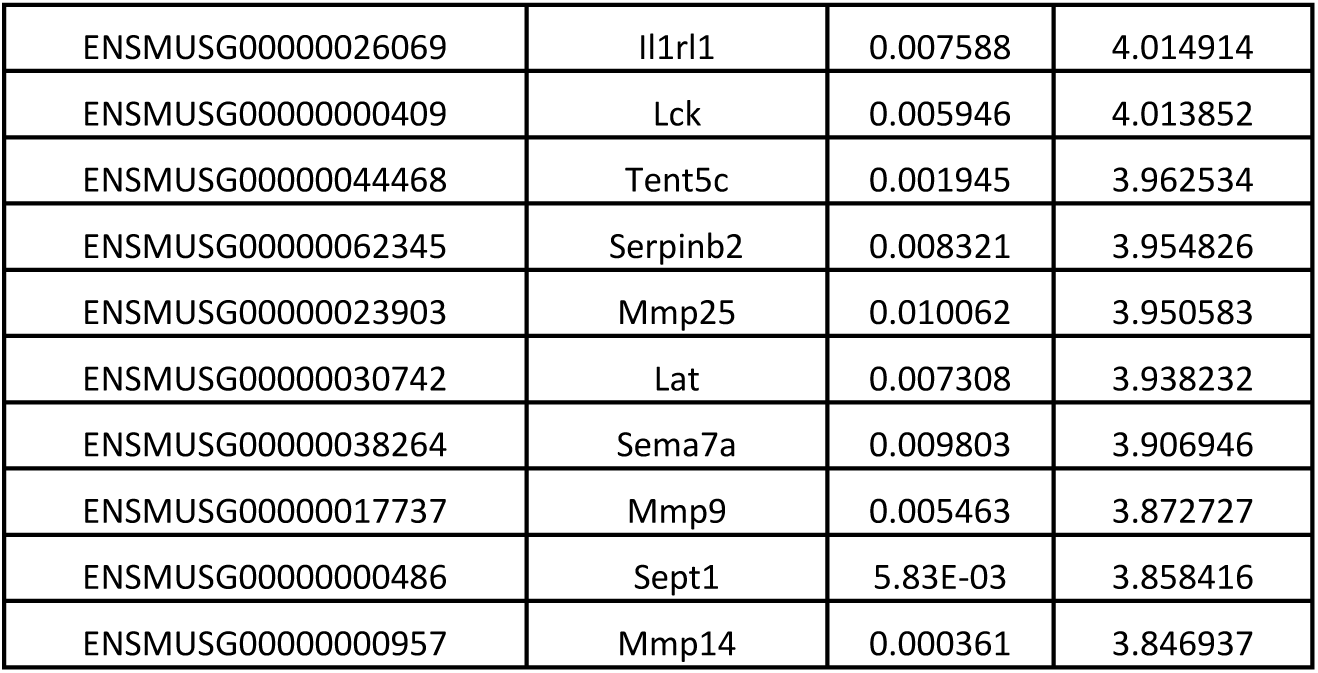
Top 50 Differentially expressed Genes upregulated in *A. fumigatus* treated mice compared to naïve controls. Positive fold change indicates increased expression in *A. fumigatus* treated mice.

**Table S2:**
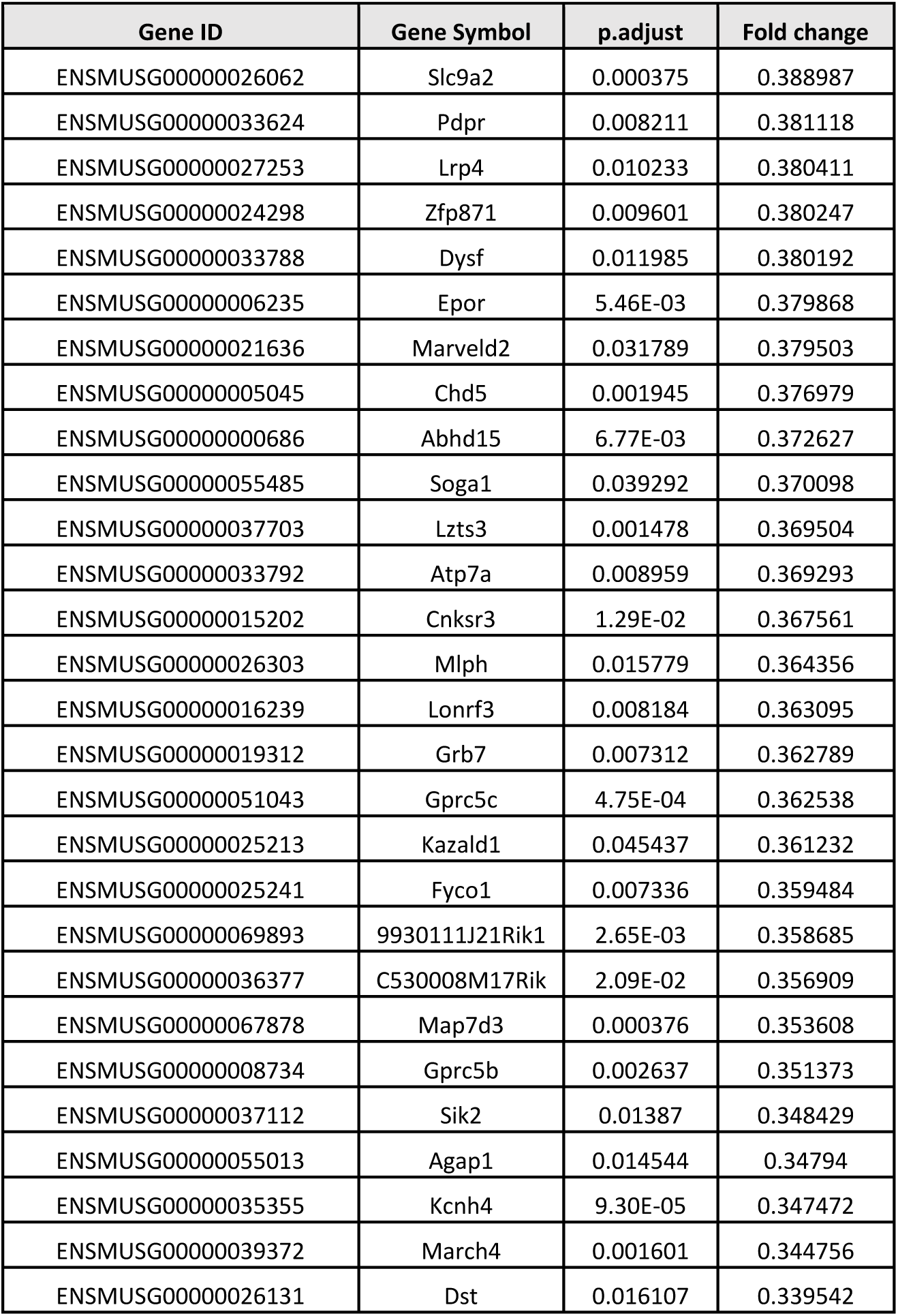
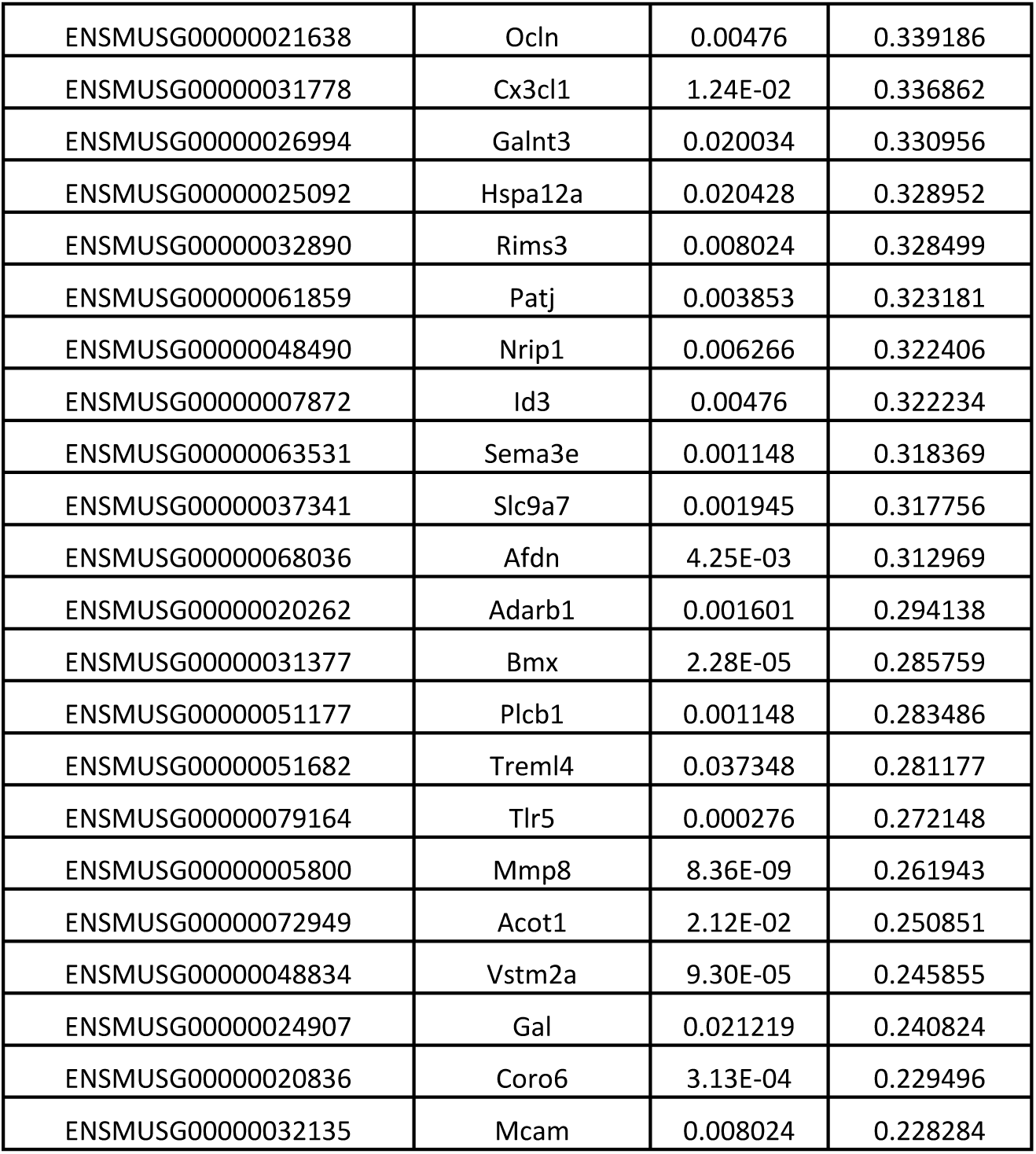
Top 50 Differentially expressed Genes downregulated in *A. fumigatus* treated mice compared to naïve controls. Positive fold change indicates increased expression in *A. fumigatus* treated mice.

**Table S3:** Top 50 Enriched GO Pathways for up and downregulated genes in *A. fumigatus* treated mice compared to naïve controls

| Upregulated Genes |  | Downregulated Genes |  |
| --- | --- | --- | --- |
| GO Pathway | p.adjust | GO Pathway | p.adjust |
| leukocyte cell-cell adhesion | 5.03E-26 | cell junction assembly | 0.006476 |
| regulation of T cell activation | 4.78E-22 | autophagy | 0.006476 |
| regulation of leukocyte cell-cell adhesion | 2.95E-21 | process utilizing autophagic mechanism | 0.006476 |
| leukocyte migration | 5.57E-19 | regulation of GTPase activity | 0.006476 |
| regulation of hemopoiesis | 3.72E-17 | axonogenesis | 0.021392 |
| T cell differentiation | 9.15E-17 | regulation of cell size | 0.021392 |
| lymphocyte differentiation | 1.94E-16 | developmental growth involved in morphogenesis | 0.021392 |
| regulation of leukocyte proliferation | 2.61E-16 | negative regulation of locomotion | 0.021392 |
| regulation of leukocyte differentiation | 5.16E-16 | regulation of nitric oxide biosynthetic process | 0.021392 |
| positive regulation of cell-cell adhesion | 5.16E-16 | regulation of nitric oxide metabolic process | 0.024836 |
| myeloid leukocyte migration | 5.16E-16 | regulation of axonogenesis | 0.028349 |
| leukocyte proliferation | 5.16E-16 | regulation of protein-containing complex disassembly | 0.028349 |
| adaptive immune response based on somatic recombination of immune receptors built from immunoglobulin superfamily domains | 9.99E-16 | nitric oxide biosynthetic process | 0.028349 |
| negative regulation of leukocyte cell-cell adhesion | 1.07E-15 | macroautophagy | 0.028349 |
| negative regulation of leukocyte activation | 1.85E-15 | positive regulation of GTPase activity | 0.028349 |
| regulation of mononuclear cell proliferation | 2.85E-15 | positive regulation of nitric oxide biosynthetic process | 0.028349 |
| mononuclear cell proliferation | 3.16E-15 | nitric oxide metabolic process | 0.028349 |
| positive regulation of cell activation | 3.16E-15 | lipid catabolic process | 0.028349 |
| positive regulation of leukocyte activation | 7.08E-15 | axon extension | 0.028349 |
| negative regulation of lymphocyte activation | 1.02E-14 | tight junction assembly | 0.028349 |
| negative regulation of cell activation | 1.31E-14 | reactive nitrogen species metabolic process | 0.028349 |
| leukocyte chemotaxis | 1.60E-14 | potassium ion transmembrane transport | 0.028349 |
| lymphocyte proliferation | 1.60E-14 | positive regulation of nitric oxide metabolic process | 0.028349 |
| regulation of lymphocyte proliferation | 1.69E-14 | neuron projection extension | 0.028672 |
| cytokine-mediated signaling pathway | 1.96E-14 | positive regulation of transmembrane transport | 0.028672 |
| negative regulation of T cell activation | 2.83E-14 | cytoskeleton-dependent intracellular transport | 0.029741 |
| mononuclear cell migration | 3.18E-14 | tight junction organization | 0.029918 |
| leukocyte mediated immunity | 3.74E-14 | central nervous system neuron development | 0.030221 |
| cell chemotaxis | 5.32E-14 | activation of GTPase activity | 0.031969 |
| negative regulation of cell adhesion | 1.06E-13 | cellular response to insulin stimulus | 0.032058 |
| CD4-positive, alpha-beta T cell activation | 1.36E-13 | regulation of autophagy | 0.032058 |
| regulation of T cell proliferation | 1.36E-13 | glycerolipid metabolic process | 0.035195 |
| negative regulation of cytokine production | 2.08E-13 | regulation of metal ion transport | 0.036004 |
| regulation of leukocyte migration | 2.35E-13 | cellular lipid catabolic process | 0.036004 |
| negative regulation of cell-cell adhesion | 3.19E-13 | regulation of synapse organization | 0.037815 |
| T cell proliferation | 3.99E-13 | negative regulation of cell migration | 0.041237 |
| granulocyte migration | 4.61E-13 | regulation of axon extension | 0.041237 |
| positive regulation of leukocyte cell-cell adhesion | 5.16E-13 | regulation of synapse structure or activity | 0.041237 |
| positive regulation of leukocyte migration | 5.44E-13 | actin cytoskeleton reorganization | 0.044417 |
| alpha-beta T cell activation | 5.74E-13 | potassium ion transport | 0.044417 |
| positive regulation of T cell activation | 7.25E-13 | transport along microtubule | 0.045424 |
| cell-substrate adhesion | 2.06E-12 | regulation of cellular component size | 0.045427 |
| negative regulation of immune response | 2.34E-12 | regulation of developmental growth | 0.045427 |
| positive regulation of leukocyte differentiation | 3.17E-12 | negative regulation of cell motility | 0.045427 |
| positive regulation of hemopoiesis | 3.17E-12 | limb development | 0.00771 |
| positive regulation of lymphocyte activation | 3.28E-12 | mitotic spindle assembly | 0.00771 |
| regulation of adaptive immune response | 5.63E-12 | protein localization to cilium | 0.008186 |
| regulation of immune effector process | 7.03E-12 | spindle assembly | 0.008186 |
| regulation of lymphocyte differentiation | 7.34E-12 | positive regulation of microtubule polymerization | 0.009335 |
| lymphocyte mediated immunity | 1.77E-11 | regulation of smoothened signaling pathway | 0.009442 |

**Table S4:**
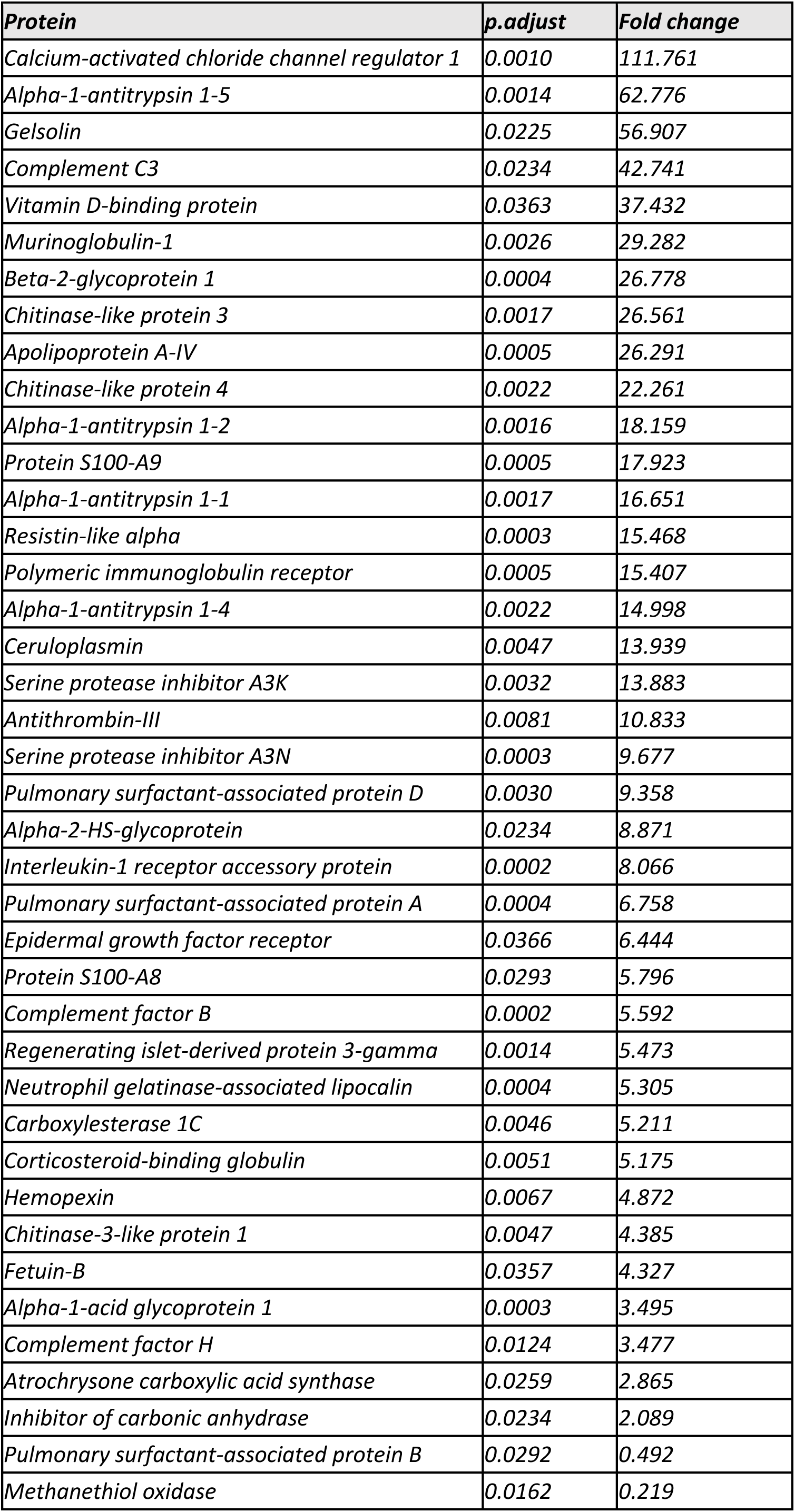
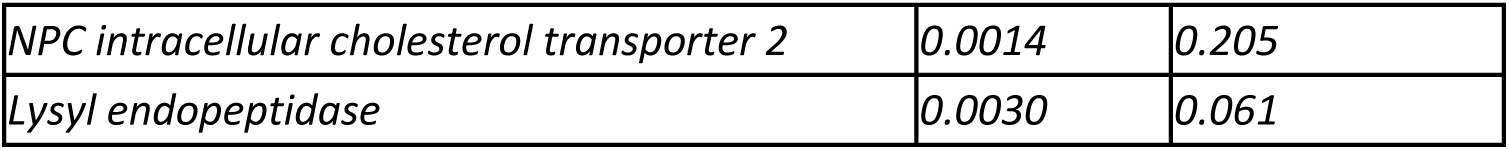
Differentially expressed proteins in *A. fumigatus* treated mice compared to naïve controls. Positive fold change indicates increased expression in *A. fumigatus* treated mice.

